# MetaMuse: A Multi-Agent AI System for Biomedical Metadata Curation and Harmonization

**DOI:** 10.64898/2026.04.12.718044

**Authors:** Ekansh Mittal, Elon Litman, Tyler Myers, Vinayak Agarwal, Ashwin Gopinath, Timothy Kassis

## Abstract

Inconsistent and unstructured metadata in public biomedical repositories, such as the Gene Expression Omnibus (GEO), severely limits data discoverability and research reproducibility. To address this, we introduce MetaMuse, a modular, multi-agent artificial intelligence framework designed to autonomously extract, validate, and standardize unstructured biomedical metadata. Operating through a three-stage architecture utilizing large language model agents, specialized CuratorAgents contextually extract candidate values for specific target metadata fields. A centralized ArbitratorAgent enforces cross-field logical consistency to prevent contradictory annotations. Finally, a NormalizerAgent leveraging a domain-specific semantic search model (SapBERT) maps these free-text candidates to formal ontological terms. We evaluated MetaMuse on a gold-standard dataset of manually curated GEO samples, achieving over 95% curation accuracy across key target metadata fields, and demonstrated robust scalability on a broader dataset of 400 samples. Notably, MetaMuse avoids data hallucination by defaulting to conservative false negatives when evidence is ambiguous, thereby preserving strict data integrity. By providing a fully auditable and context-aware curation pipeline, MetaMuse offers a scalable solution for enriching public data repositories and accelerating reproducible, data-driven scientific discovery.

## 1 Introduction

### Motivation

Reproducibility and data discoverability are pressing concerns in biomedical research; over 80% of scientists perceive a “reproducibility crisis” (Baker, 2016). A major contributing factor is the highly variable quality of dataset annotations, which makes it difficult for independent teams to replicate experiments or reuse data (Gonçalves and Musen, 2019; Leipzig et al., 2021). Consequently, detailed metadata integrity is now recognized as a fundamental determinant of research credibility, providing the necessary provenance to validate data-driven findings (Caliskan et al., 2023a,b).

High-quality metadata also makes data findable and usable, aligning directly with the FAIR data principles (Wilkinson et al., 2016; Hughes et al., 2023). Uniformly capturing experimental descriptors facilitates downstream applications like machine learning; conversely, critical information buried in unstructured text makes systematic querying nearly impossible (Wang et al., 2017; Teng et al., 2018). Ultimately, tagging experiments with controlled ontological terms enables large-scale, cross-study interoperability, unlocking analyses that would otherwise be infeasible (Mungall et al., 2012; Hoyt et al., 2022; Pouran Ben Veyseh et al., 2022).

Unfortunately, much of today’s public biomedical data lacks the rigorous, structured annotation required for modern computational analyses. In the Gene Expression Omnibus (GEO), for example, the absence of strictly enforced upload schemas means that critical experimental descriptors are frequently relegated to free-form text fields by data submitters (Wang et al., 2017; Teng et al., 2018). This leads to highly variable annotation quality across different studies (Gonçalves and Musen, 2019), a systemic issue that crowdsourcing initiatives have only just begun to quantify (Zaveri et al., 2019). As a result, essential biological attributes—such as tissue origin or primary disease state—remain effectively “hidden in plain sight.” They exist within narrative descriptions but completely defy systematic search or cross-study comparison. This heterogeneity is stark even in basic fields; for instance, simple binary variables like donor sex are frequently encoded using wildly inconsistent conventions (e.g., “M”, “male”, “1”) (Wang et al., 2017; Gonçalves and Musen, 2019). Gonçalves and Musen (2019) documented such anomalies across more than 11 million sample records, noting that they severely “impede search and secondary use.” Ultimately, this widespread reliance on cursory, unstructured metadata directly fuels the dual crises of reproducibility and discoverability, keeping valuable datasets functionally invisible to automated retrieval systems (Hughes et al., 2023; Caliskan et al., 2023b).

### Objectives and Contributions

The primary objective of this research is to resolve this critical bottleneck of unstructured metadata within public repositories like GEO. We aim to overcome the historical trade-off between the high accuracy of labor-intensive manual curation and the scalability, but lower reliability, of automated heuristic pipelines. To achieve this, we developed MetaMuse, an automated framework designed to parse, validate, and standardize complex biomedical metadata into formal ontological terms while maintaining strict experimental context and data integrity.

This paper presents the following key contributions:

- **Multi-Agent AI Architecture:** We introduce MetaMuse, a modular system utilizing distinct specialized agents, Curator, Arbitrator, and Normalizer, to process raw dataset inputs into structured, standardized formats.
- **Context-Aware Extraction:** We deploy CuratorAgents capable of parsing literature and dataset metadata to extract contextually relevant candidate values, utilizing a conservative approach that significantly minimizes false positive hallucinations.
- **Cross-Field Logical Arbitration:** We implemented an ArbitratorAgent that evaluates metadata holistically, enforcing logical consistency across fields (e.g., preventing contradictory disease and cell line pairings) through an iterative self-correction loop.
- **High-Fidelity Normalization:** We utilize a semantic search module powered by SapBERT to accurately map colloquial and highly variable biomedical terms to standardized ontology identifiers, such as MONDO and UBERON.
- **Auditable & Scalable Validation:** We empirically demonstrate >95% curation accuracy across key fields on a gold-standard GEO dataset. Furthermore, MetaMuse generates a fully transparent, auditable trail for every programmatic decision, directly supporting the principles of reproducible science. The remainder of this paper reviews prior curation efforts (Section 2), details the multi-agent MetaMuse architecture (Section 3), evaluates its performance on real-world GEO datasets (Section 4), and discusses the ongoing challenges of ontology normalization (Section 5).

## 2 Related Work

Extensive efforts have been made to address the biomedical metadata bottleneck, ranging from crowdsourcing initiatives to advanced language models; however, existing solutions frequently struggle to balance quality, scalability, and auditability.

### The Accuracy vs. Scalability Trade-off

Early efforts to improve biomedical metadata highlight a fundamental trade-off between annotation accuracy and scalability. On one end of the spectrum, human-in-the-loop manual curation remains the gold standard for precision. Dedicated projects, such as GEOMetaCuration, employ domain experts to painstakingly correct errors and align entries with formal ontologies (Jiao et al., 2018). However, manual review is prohibitively slow and fundamentally non-scalable, rendering it poorly suited for generating the massive training datasets required by modern machine learning applications (Elucidata, 2025; Garcia et al., 2025). Even distributed crowdsourcing initiatives cannot feasibly cover the breadth of metadata across the expanding biomedical landscape (Zaveri et al., 2019; Coe et al., 2017). To circumvent these labor bottlenecks, researchers initially deployed automated text-mining pipelines. First-generation systems relied heavily on rule-based heuristics, utilizing regular expressions and keyword scanners to identify simple variables like age or donor sex (Wang et al., 2017; Teng et al., 2018). While these systems achieved the necessary throughput, they sacrificed accuracy and generalization, frequently failing when confronted with evolving metadata schemas (Kibbe et al., 2010).

### Challenges in Context and Cross-Field Consistency

Beyond basic extraction, reliable metadata curation demands deep contextual reasoning and cross-field consistency—requirements that expose the limits of traditional NLP. Trivial string-matching algorithms lack the domain awareness necessary to resolve ambiguities, famously misinterpreting the species identifier *Mus musculus* as an indicator of muscle tissue (Hawkins et al., 2022). Furthermore, biological descriptors are heavily interdependent; for instance, a sample’s annotated disease state cannot logically contradict its cell line. Ontology-driven tools like MetaSRA improved upon naïve taggers by mapping free text to controlled vocabularies, but they still struggled with complex repository data, frequently producing false positives that necessitated manual intervention (Hawkins et al., 2022). While modern multi-task NLP frameworks like BiomedCurator show promise, their architectures are often optimized for structured literature (e.g., PubMed abstracts) rather than addressing the severe sample-level heterogeneity and cross-field constraint violations found in unstructured repositories (Sohrab et al., 2022).

### Gaps in Auditability and Maintainability

Finally, the long-term adoption of any curation framework depends heavily on its auditability and maintainability. Earlier algorithmic pipelines were often built as monolithic, one-off solutions that suffered from cumbersome runtimes and reliance on quickly obsolete software infrastructure (Hawkins et al., 2022). While recent attention has enthusiastically shifted toward large language models (LLMs). Recent pipelines have utilized LLMs for sequencing repositories (Li et al., 2023), while others have specifically targeted the annotation of GEO studies (Kaczmarek et al., 2025; Zhang et al., 2025). Current implementations still exhibit significant evaluation gaps. For example, recent industry-led multi-agent systems report strong reliability across diverse fields, yet they frequently omit transparent, head-to-head benchmarking on public corpora (Elucidata, 2025). Critically, they lack span-level provenance, per-field error decomposition, and explicit governance rules for cross-field arbitration. Without auditable evidence trails and modular designs that allow researchers to swap out mapping ontologies without retraining (Hoyt et al., 2022; Pouran Ben Veyseh et al., 2022), users cannot independently verify the integrity of the data. Overall, the field still lacks a repository-focused LLM framework that pairs high-fidelity curation with transparent, reproducible provenance.

## 3 Methods

MetaMuse has a modular, three-stage architecture. It has been built specifically to process the raw unstructured metadata found in the Gene Expression Omnibus (GEO), but can be adapted to handle raw metadata from any source. In the first stage, for given sample IDs, MetaMuse pulls the relevant sample (GSM), series (GSE), and abstract metadata, removes irrelevant fields, and packages the data from all sources into a single unit, establishing the evidence boundary for the run. Second, preprocessing performs a fast pass to determine the sample type (whether it is from a primary sample or cell line study) and to organize work into fixed-size batches by type. Third, conditional processing performs field-specific curation, an optional arbitration-driven correction pass, and finally normalization to controlled vocabularies.

MetaMuse currently supports the following target metadata fields and respective ontologies:

**Table 1:** Currently supported metadata fields and respective ontologies.

| *Metadata Field* | *Ontology* | *Version* | *Relevant Sample Type* |
| --- | --- | --- | --- |
| Disease | MONDO | 2024-08-06 | All Samples |
| Tissue | UBERON | 2024-08-07 | Primary Samples |
| Treatment | ChEMBL | CHEMBL35 | All Samples |
| Organism | NCBI Taxonomy | 2023-06-20 | All Samples |
| Ethnicity | HANCESTRO | 3.0 | Primary Samples |
| Developmental Stage | HSAPDV | 2024-05-28 | Primary Samples |
| Gender | PATO | 2023-05-18 | Primary Samples |
| Cell Line | CLO | 2.0 (2025-06-01) | Cell Lines |
| Sample Type | Primary Sample vs Cell Line | NA | All Samples |
| Age | NA | NA | Primary Samples |
| PubMed ID | NA | NA | All Samples |
| Instrument ID | NA | NA | All Samples |

**Figure 1:**
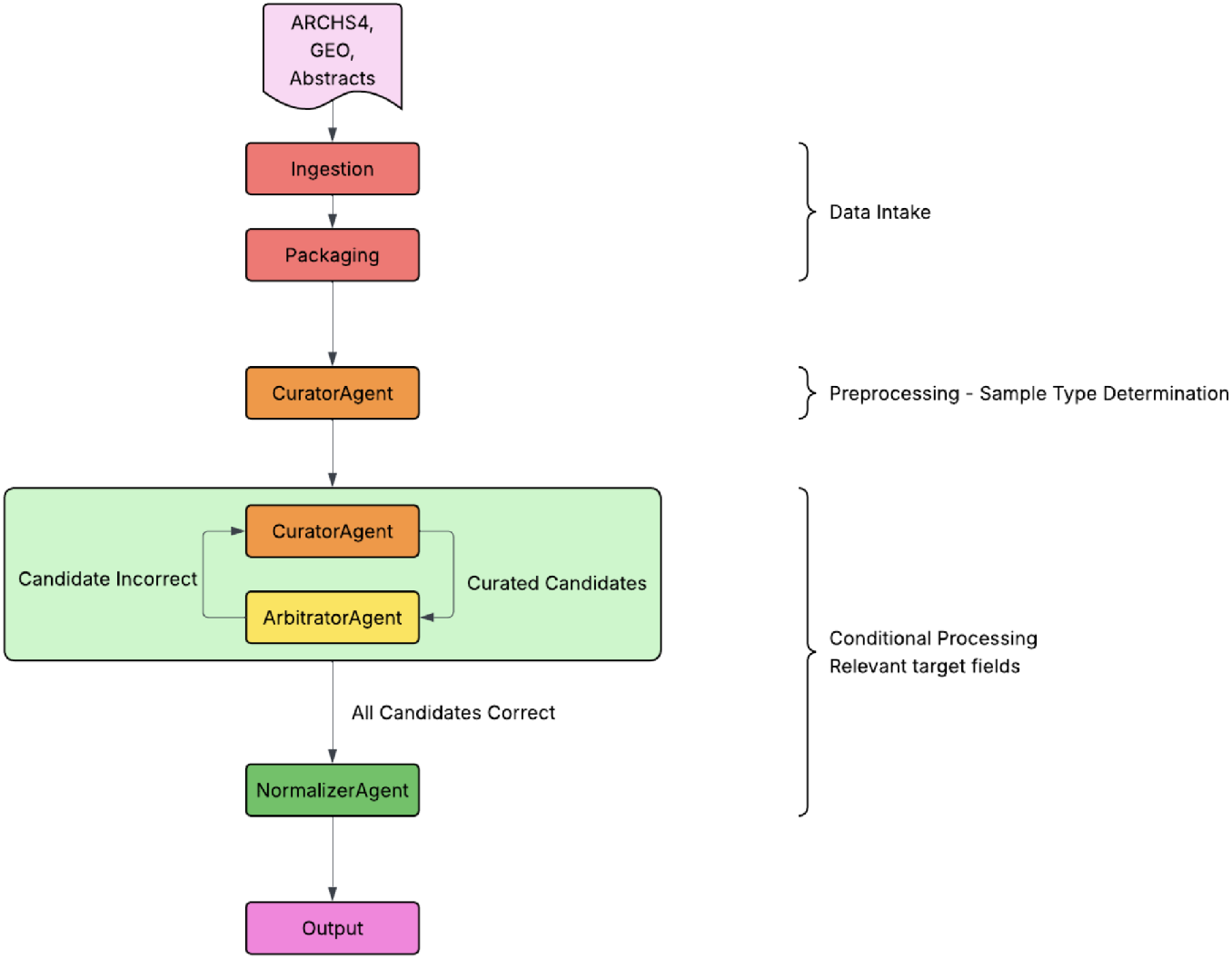
MetaMuse Architecture

While the initial data intake is deterministic, the subsequent preprocessing and conditional processing stages rely on an ensemble of independent large language model (LLM) agents, built upon the `openai/agents-sdk` (OpenAI, 2024). To optimize for both speed and complex reasoning, MetaMuse routes tasks to different models based on computational difficulty. Table 2 summarizes the specialized agents deployed in the pipeline, their designated models, and their core responsibilities.

**Table 2:** Overview of MetaMuse agents, their underlying language models, and their primary functions within the pipeline.

| *Agent* | *Model Version* | *Primary Purpose* |
| --- | --- | --- |
| CURATORAGENT (Preprocessing) | Gemini-2.5-flash | Determines sample type (primary vs. cell line) to organize downstream tasks. |
| CURATORAGENT (Curation) | Gemini-2.5-pro | Extracts contextual candidate values for target metadata fields from raw text. |
| ARBITRATORAGENT | Gemini-2.5-pro | Reviews cross-field consistency, identifies errors, and provides correction feedback. |
| NORMALIZERAGENT | Gemini-2.5-flash | Maps curated free-text candidates to formal ontological terms via semantic search. |

### 3.1 Data Intake

As described previously, the data intake stage is responsible for ingesting data from relevant sources, such as GEO and PubMed, and preparing it for processing by the downstream agents. Originally, the NCBI Entrez API was used to source the data, but the rate limits became prohibitively slow, so two SQLITE databases were prepared/sourced for GEO and PubMed. These were prefiltered to only include the relevant GSM (sample), GSE (series), and PubMed (abstract) IDs. Once all the relevant metadata has been ingested from the two SQLITE databases, extraneous fields, such as contact information, addresses, and author names, are discarded. All relevant data for all samples in the query is then packaged into a JSON dump to allow easy access for downstream agents.

### 3.2 Preprocessing

The preprocessing stage determines whether a sample/study is sourced from a primary sample (a patient) or a cell line. Making this determination is crucial because it vastly changes the target metadata fields that will be of interest for a sample. Particularly, for primary samples, patient demographics (age, ethnicity, developmental stage, gender) and source tissue information are critically important, whereas they are not available/relevant for cell line samples. On the other hand, there is no point in searching for cell line identifiers in primary sample studies. Additionally, the scope/context of some target metadata fields, such as treatment, changes depending on the sample type. Making this sample type determination before actually curating any fields ensures that the extracted information is relevant and taken into context. The Preprocessing stage consists of one agent: CuratorAgent, (described in 3.3) restricted to the sample_type field. The end result of the preprocessing stage is a mapping between samples and sample types. This helps inform the conditional processing stage during sample batching.

### 3.3 Conditional Processing

The Conditional Processing stage is responsible for curating and normalizing all remaining target metadata fields. To achieve this, the system instantiates an ensemble of approximately ten CuratorAgents per sample—dedicating one specific agent to each target metadata field (e.g., disease, organ, tissue). This decentralized approach not only improved performance in our initial experiments by allowing each agent to act as a focused expert, but it also enabled efficient batching of multiple samples into a single task without sacrificing accuracy.

#### CuratorAgent

Each CuratorAgent reads the raw JSON from the data intake stage and executes a multi-step reasoning process to extract candidate values. It first scans all three text sources for potential candidates before settling on a final extraction. What distinguishes the CuratorAgent is that it is strictly context-aware and fully auditable. It understands that some information sourced from series and abstract-level metadata may not be relevant to the specific sample in question. For example, if “Breast Cancer” is mentioned in a study’s abstract merely as a potential future application of the research, the CuratorAgent will recognize that context and correctly decline to report it as the sample’s disease state. Ultimately, the agent outputs the final candidate alongside the context in which it was found, the rationale for its selection, and its confidence score.

Once all target metadata fields for a sample have been successfully processed from raw metadata into curated candidates, the pipeline invokes the next component: the ArbitratorAgent.

#### ArbitratorAgent

This agent’s responsibility is to review the CuratorAgent’s results and provide feedback to it. While the CuratorAgent is only focused on a single target metadata field, the ArbitratorAgent reviews all target metadata fields for a given sample and checks the CuratorAgent’s results against the original source metadata. It is important to consider all target metadata fields together to prevent conflicts in information. If the cell line CuratorAgent identifies “MDA-MB-231” (Triple negative breast cancer cell line), and the disease CuratorAgent identifies something other than Breast Cancer, the ArbitratorAgent will pick up on this error. The ArbitratorAgent returns three values per target metadata field: 1) a binary value of whether each specific target metadata field is correct, 2) the reason the value is incorrect, and 3) the suggested new value if it was incorrect. The workflow parses this output and reinvokes CuratorAgents to make corrections for specific target metadata fields. This loop continues until the ArbitratorAgent deems all target metadata fields correct, or until a maximum of 3 iterations is reached, at which point the sample is flagged for manual review.

Once all target metadata fields have been curated and deemed correct by the ArbitratorAgent, the curated candidate values are passed to the final agent, NormalizerAgent.

#### NormalizerAgent

This agent’s responsibility is to map all curated free-text candidate values to their corresponding normalized ontological terms. For example, the colloquial term “Breast Cancer” is accurately mapped to its formal ID, “MONDO:0007254.” To achieve this, the NormalizerAgent leverages a semantic search module that evaluates the free-text candidate against standard ontology dictionaries. This semantic approach ensures that results are deterministic and that highly variable synonyms are reliably linked to the correct concept. The search yields a short, ranked list of the top candidate matches (*n* = 5) along with confidence scores. The NormalizerAgent then analyzes these candidates and returns the best ontological match, outputting the final ID alongside the reasoning for its selection. Complete implementation details of the semantic search indexing and retrieval model are provided in Appendix 6.

Once the NormalizerAgent completes, all the curated and normalized values are saved to a single CSV or Parquet file. All of the raw CuratorAgent, ArbitratorAgent, and NormalizerAgent JSON outputs are stored for auditability. This approach provides a minimal yet robust method for converting unstructured biomedical metadata into well-defined structured fields.

## 4 Results

To validate the performance of MetaMuse, we conducted a series of evaluations designed to measure its accuracy, scalability, and the fidelity of its ontology normalization. Our findings demonstrate that MetaMuse achieves high accuracy on a manually curated gold-standard dataset and maintains strong performance at scale. Furthermore, the system’s outputs are context-aware and fully auditable, addressing key limitations of previous metadata curation approaches.

### 4.1 High Curation Accuracy on a Gold-Standard Validation Set

We first benchmarked MetaMuse against a manually curated “ground truth” of 100 randomly selected GEO samples, selected to represent a diverse set of primary tissue and cell line studies. The system demonstrated high accuracy across the majority of key metadata fields, proving its ability to handle complex, real-world metadata. As shown in Figure 2, per-field accuracy was exceptionally strong for all fields, as shown by accuracies > 95% for all fields.

**Figure 2:**
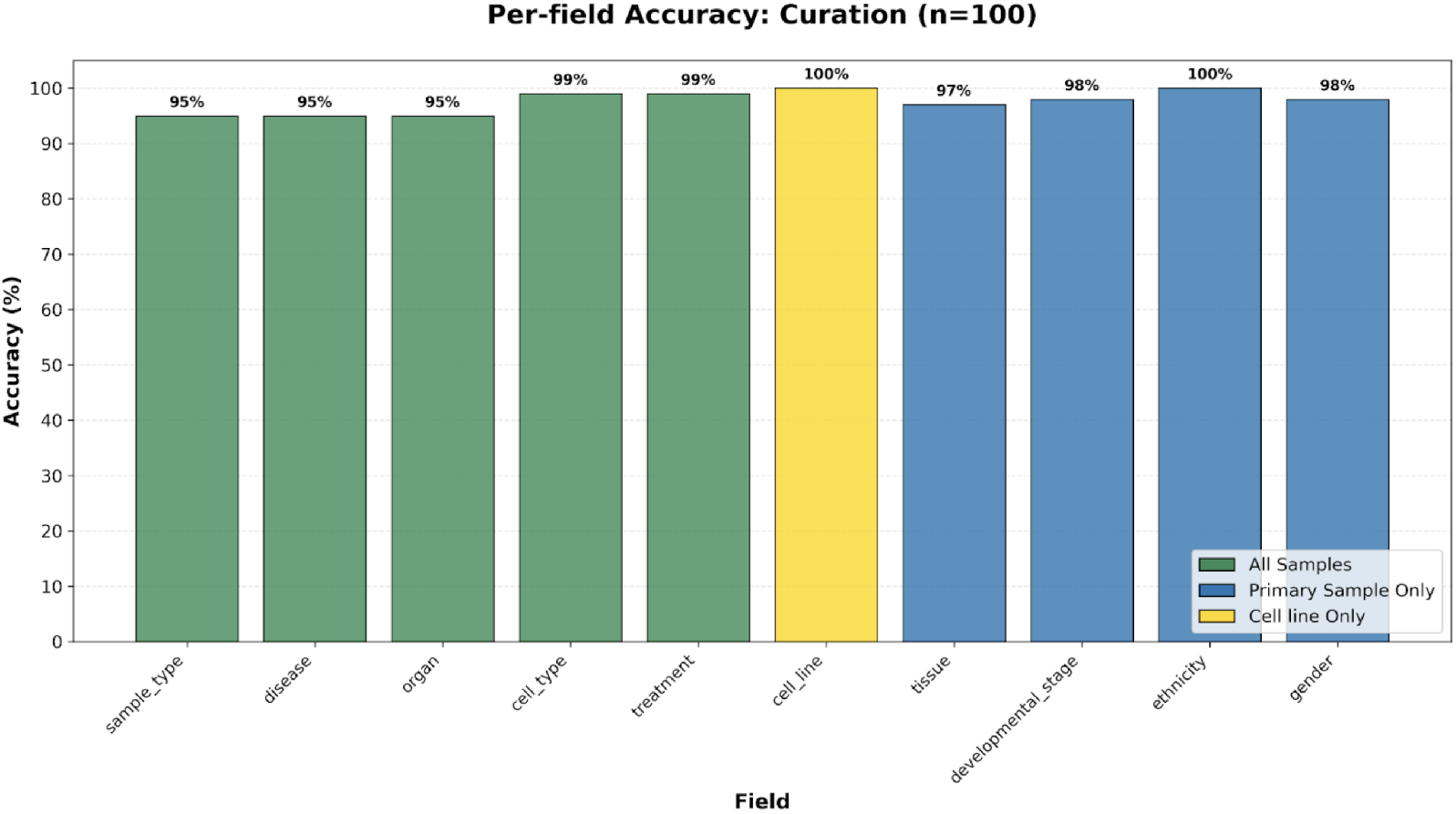
Per-field curation (n=100) accuracy on a manually curated set of 100 samples.

A manual audit of the discrepancies revealed that the predominant error type was false negatives (i.e., the CuratorAgent conservatively reported “None reported” when a value was present in the source text) rather than false positives (hallucinations) (Table 3). For instance, in the tissue field (86% accuracy), several errors were due to the agent missing a subtle mention of the tissue type embedded deep within a lengthy abstract or in a non-standard characteristics field. This conservative behavior is a deliberate design choice; it indicates that the system is highly precise and will omit information rather than invent it. In biomedical research, where an incorrect data point can invalidate an entire analysis, this refusal to hallucinate is a critical feature in generating trustworthy, reliable metadata.

**Table 3:**
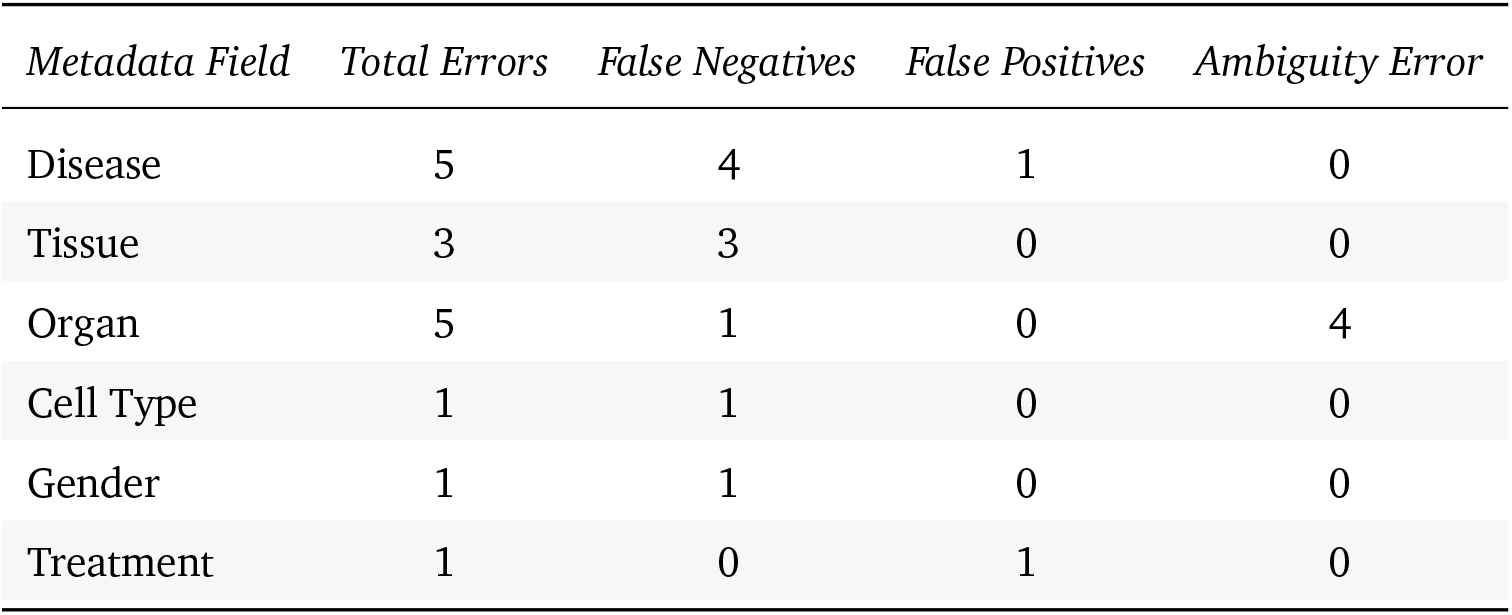
Error distribution by type for supported metadata fields (n=100 samples).

| *Metadata Field* | *Total Errors* | *False Negatives* | *False Positives* | *Ambiguity Error* |
| --- | --- | --- | --- | --- |
| Disease | 5 | 4 | 1 | 0 |
| Tissue | 3 | 3 | 0 | 0 |
| Organ | 5 | 1 | 0 | 4 |
| Cell Type | 1 | 1 | 0 | 0 |
| Gender | 1 | 1 | 0 | 0 |
| Treatment | 1 | 0 | 1 | 0 |

### 4.2 Scalable Performance and High-Fidelity Normalization

To evaluate the scalability and broader performance of MetaMuse, we processed an additional 400 samples and used an external LLM (Gemini 2.5 pro) evaluator to score the results. As shown in Figure 3 the system’s curation accuracy remained high across this larger, more diverse dataset, consistent with the performance observed on the manual validation set and demonstrating its capability to handle high-throughput curation tasks.

**Figure 3:**
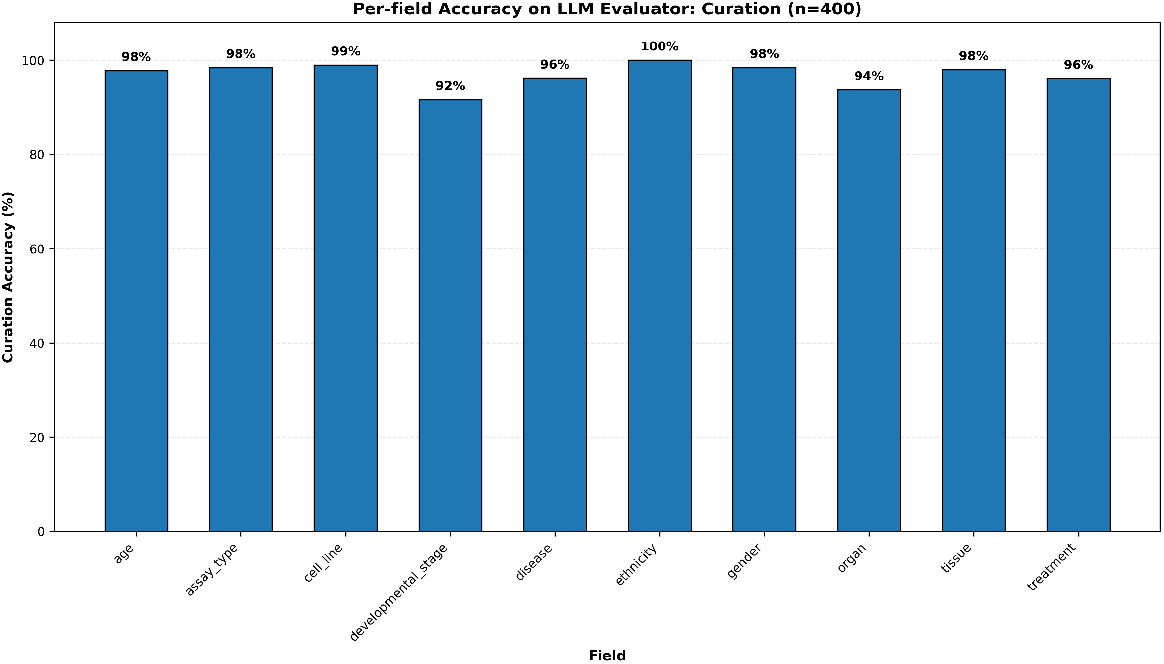
Per field curation accuracy of MetaMuse on 400 samples, evaluated by Gemini 2.5 pro.

### 4.3 Normalization

A critical step in our pipeline is the normalization of curated free-text values to controlled ontology terms. We assessed the fidelity of this step by comparing accuracy scores before and after normalization by the NormalizerAgent. As shown in Figure 4, our results reveal a clear trend: while context-aware curation achieves exceptionally high accuracy, formal ontology normalization remains the primary bottleneck in the pipeline.

**Figure 4:**
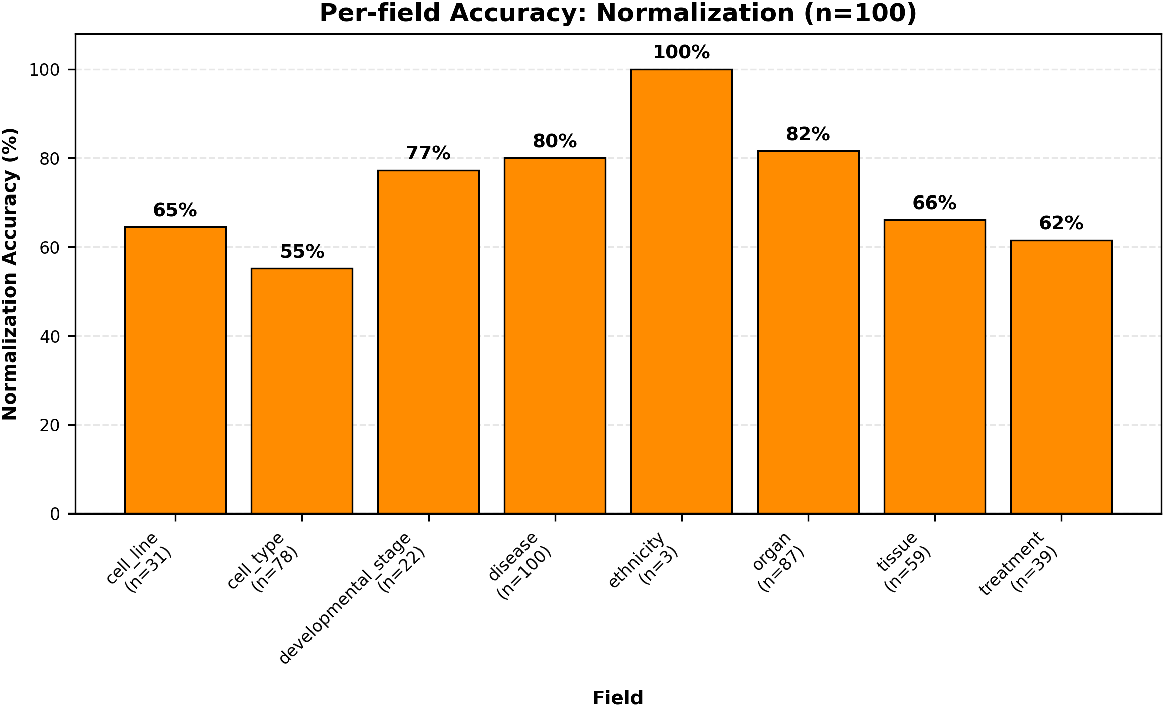
Per-field normalization (n=100) accuracy on a manually curated set of 100 samples.

For instance, accuracy dropped significantly post-normalization in fields like disease (95% to 80%) and tissue (97% to 66%). While the underlying SapBERT model successfully bridges the gap between many colloquial terms (e.g., “mammary tumor”) and formal ontology concepts (“breast carcinoma,” MONDO:0007254), it frequently struggles with highly specialized or composite entities. This limitation is most evident in the sharp accuracy drops for cell type (55%) and treatment (62%). A common failure mode occurs when SapBERT encounters high-granularity text; for example, it may erroneously map a highly specific descriptor like “CD8+ effector memory T cell” to a broader parent node (e.g., “T cell”), which is penalized as an error under strict validation constraints. Similarly, combinatorial treatment descriptions often lose crucial context during the embedding process. Ultimately, these results highlight that standardizing highly heterogeneous biomedical text into rigid ontological frameworks remains a fundamental challenge, even when the initial data extraction is nearly flawless.

## 5 Discussion

In this work, we introduced MetaMuse, a multi-agent AI framework that produces high-quality, ontology-aligned biomedical metadata at scale. By ensuring cross-field consistency and maintaining strict context, MetaMuse represents a significant step forward in making public data repositories, such as GEO, more FAIR (Findable, Accessible, Interoperable, and Reusable).

A key innovation driving this performance is the ArbitratorAgent, which directly addresses a critical failure point in prior automated systems: the inability to validate information across different target metadata fields. While simple NLP pipelines can extract individual terms, they often fail to recognize logical impossibilities, such as a sample being annotated with a lung cancer-specific cell line but a disease state of liver cancer. The iterative correction loop between our expert CuratorAgents and the ArbitratorAgent mimics the consensus-driven process of human expert curation, allowing the system to self-correct and produce a coherent, biologically plausible final record. This arbitration is the primary driver of the high curation accuracy reported in our results.

Furthermore, our evaluation highlighted the system’s high precision and conservative nature. The finding that most curation errors were false negatives (“missed cues”) rather than false positives (“hallucinations”) is a crucial and deliberate design outcome. In data-driven science, the introduction of erroneous data is far more damaging than the omission of a data point. By prioritizing accuracy and avoiding the invention of information, METAMUSE produces a reliable substrate for downstream applications where data integrity is paramount.

Beyond raw accuracy and data integrity, a core operational advantage of MetaMuse is its capacity for context-aware curation coupled with full auditability. To this end, 100% of the outputs generated during our 400-sample evaluation successfully included a traceable rationale chain and JSON evidence log. The system demonstrated a quantitative capacity for nuanced understanding, such as correctly synthesizing sample-level details with series-level context to identify control samples (e.g., mapping an uninfected macrophage in a tuberculosis study to “control [tuberculosis],” as shown in Figure 5).

**Figure 5:**
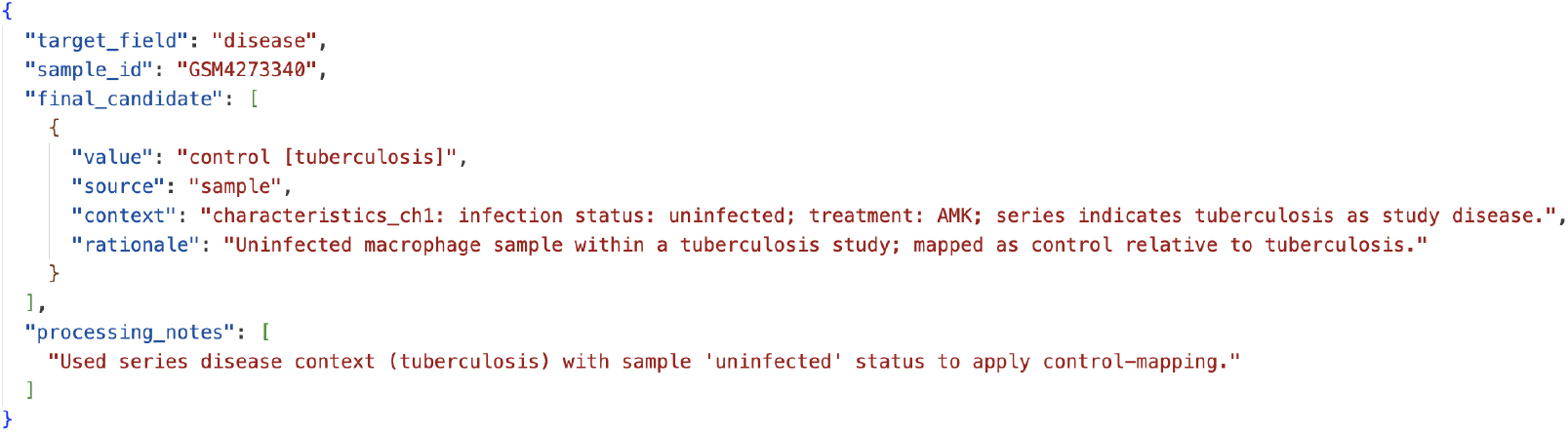
Example of curation output on a sample, displaying control mapping, as well as MetaMuse’s source’s and rationale

The rationale provided in these outputs creates a transparent trail for every curated value. This allows a human reviewer to verify the complete decision-making process: from the initial evidence fragments extracted by the CuratorAgent, through the cross-field consistency checks performed by the ArbitratorAgent, and concluding with the final ontology mapping from the NormalizerAgent.

This end-to-end transparency is essential for building trust in automated systems and directly supports the overarching goal of improving research reproducibility.

However, our results also reveal a notable limitation: formal ontology normalization currently serves as the pipeline’s primary bottleneck. As observed in Section 4.3, while MetaMuse’s initial curation is highly accurate, fidelity drops noticeably when mapping those curated free-text values to strict ontology IDs. The success of the SapBERT-based semantic search proves that domain-specific models are necessary, yet SapBERT still struggles with highly specialized, granular, or combinatorial descriptions (such as complex cell subtypes or drug treatments). Addressing this normalization gap is a key priority for future work. Potential paths forward include fine-tuning SapBERT on a more specific corpus of high-granularity GEO annotations, or deploying an LLM-based normalizer to act as a fallback reasoning mechanism when semantic search yields low confidence scores.

Finally, MetaMuse is inherently dependent on the quality and availability of the source metadata. In cases where a GEO entry is sparsely annotated and has no linked publication, the system’s ability to curate accurately is fundamentally limited. Additionally, while the system handles most common patterns, highly unusual or complex experimental designs still require deep, domain-specific human inference. Future iterations could incorporate more sophisticated reasoning modules to automatically identify and flag these ambiguous edge cases for expert human review.

## 6 Conclusion

The challenge of curating unstructured metadata is a major bottleneck impeding progress in biomedical research and contributing to the ongoing reproducibility crisis. Our work demonstrates that a multi-agent AI system, MetaMuse, can automate this process with a high degree of accuracy and reliability. By combining specialized agents for extraction, arbitration, and normalization, our system produces auditable, context-aware, and ontology-aligned metadata at a scale unattainable by manual methods. MetaMuse provides a robust and scalable solution for enriching public data repositories, thereby improving the interoperability of these resources for reproducible scientific discovery.

## Appendix A

**Semantic Search Implementation for Normalization**

To perform the high-fidelity ontology normalization described in Section 3, the NormalizerAgent utilizes a specialized semantic search architecture. The tool embeds both the extracted free-text candidate values and the target ontology dictionary terms using the pre-trained Hugging Face model `cambridgeltl/SapBERT-from-PubMedBERT-fulltext`, a model specifically optimized for biomedical entity linking.

The resulting embeddings are stored and queried using a FAISS (Facebook AI Similarity Search) vector index, which is GPU-accelerated when hardware permits. At runtime, the system encodes the query text, normalizes the resulting vectors, and performs a *k*-nearest neighbor search over the indexed ontology terms. Results are filtered by a minimum cosine similarity threshold, and the top *n* = 5 matches are returned to the NormalizerAgent along with their similarity scores. This decoupled infrastructure ensures rapid, scalable, and highly accurate semantic retrieval even across massive biomedical ontologies.

